# Dopamine and temporal discounting: revisiting pharmacology and individual differences

**DOI:** 10.1101/2024.08.28.610170

**Authors:** Elke Smith, Hendrik Theis, Thilo van Eimeren, Kilian Knauth, Deniz Tuzsus, Lei Zhang, David Mathar, Jan Peters

## Abstract

Disorders characterised by changes in dopamine (DA) neurotransmission are often linked to changes in the temporal discounting of future rewards. Likewise, pharmacological manipulations of DA neurotransmission in healthy individuals modulate temporal discounting, but there is considerable variability in the directionality of reported pharmacological effects, as enhancements and reductions of DA signalling have been linked to both increases and reductions of temporal discounting. This may be due to meaningful individual differences in drug effects and/or false positive findings in small samples. To resolve these inconsistencies, we 1) revisited pharmacological effects of the DA precursor L-DOPA on temporal discounting in a large sample of N = 76 healthy participants (n = 44 male) and 2) examined several putative proxy measures for DA to revisit the role of individual differences in a randomised, double-blind placebo-controlled pre-registered study (https://osf.io/a4k9j/). Replicating previous findings, higher rewards were discounted less (magnitude effect). Computational modelling using hierarchical Bayesian parameter estimation confirmed that the data in both drug conditions were best accounted for by a non-linear temporal discounting drift diffusion model. In line with recent animal and human work, L-DOPA reliably reduced the discount rate with a small effect size, challenging earlier findings in substantially smaller samples. We found no credible evidence for linear or quadratic effects of putative DA proxy measures on model parameters, calling into question the role of these measures in accounting for individual differences in DA drug effects.

## 1 INTRODUCTION

We are constantly confronted with decisions between options that have an immediate benefit, and options that have a comparatively greater benefit, which, however, only materialises in the future. Individuals commonly devaluate future rewards to some extent, a phenomenon known as temporal discounting^1,2^, which may lead them to forgo future larger rewards. A more pronounceSd tendency to choose smaller immediate rewards (impulsive choice) is observed in individuals suffering from substance-use-disorders (SUDs) and behavioural addictions^3,4^. One prominent hypothesis assumes changes in dopamine (DA) neurotransmission as functional correlate of maladaptive changes in decision-making^5,6^. This is supported by a large body of human and animal evidence suggesting that DA plays a central role in reward valuation, prediction and decision-making^7–10^. Reinforcers act upon frontal and striatal brain regions innervated by dopaminergic neurons and implicated in reward processing^11–13^. Individual differences in DA D2 receptor availability in the midbrain correlate with reward valuation^14^, whereas highly impulsive individuals and individuals suffering from SUDs tend to exhibit decreased DA function (for reviews, see^6,15^).

Several studies have investigated the effects of manipulating DA neurotransmission on temporal discounting, but yielded highly mixed results. A highly-cited earlier study reported an increase in temporal discounting, i.e. stronger devaluation of future rewards, following enhancement of DA neurotransmission via the DA precursor L-DOPA^16,^. In contrast, other studies reported decreases^17–20^, no effects^21–23^, or moderating effects of impulsivity and body weight on DAergic drug effects^24^. Studies attenuating DA neurotransmission also provided mixed results, some of which reported no effects^25^, a reduction in temporal discounting^26,27^, or a genotype interaction with DAergic drug effects^28^. Some results should be interpreted with caution due to low sample sizes (e.g., *N* = 10^22^; *N* = 13^16^; *N* = 14^26^), low trial numbers (e.g. 27 intertemporal decisions^26^), or delays being confounded with probabilities^26^. Furthermore, the studies differ in the substances used to increase (e.g. L-DOPA^16^, d-amphetamine^21,29^, buproprion^21^, pramipexole^22^, tolcapone^20^, PF-06412562^23^, and haloperidol^18,19^, a DA receptor antagonist, which, at lower doses, is assumed to increase striatal DA) or decrease (e.g. amisulpride^27^, metoclopramide^26^, and acute phenylalanine/tyrosine depletion^28^) DAergic neurotransmission thereby impacting on different DAergic signalling mechanisms, such as inhibiting COMT, preventing DA reuptake, or activating different DA receptor subtypes. Since activation of D1 and D2 receptors is thought to exert differential effects on value-based decision making^30^, this variability may influence the consistency of the findings.

Another explanation for the observed inconsistencies in DA drug effects may be related to interindividual differences in baseline DA function. Specifically, it has been proposed that the relationship between DA levels and cognitive function follows an inverted U-shaped function^31^. In this model, pharmacological enhancement of DA signalling would result in either worse or better cognitive functioning, depending on an individual’s baseline DA level, a relationship that was proposed for working memory (WM) and cognitive control^31^, sensation seeking^32^, and impulsivity^24^. Individuals with higher striatal DA synthesis capacity were also more motivated to exert effort, while those with lower synthesis capacity showed a greater response to methylphenidate and sulpiride^33^. While this observation is consistent with the inverted U-shaped hypothesis, it is important to note that, despite frequent proposals of an inverted U-shaped relationship between DA levels and cognitive function, this hypothesis has rarely been directly tested by measuring DA receptor availability or synthesis capacity in individuals scoring lower and higher on cognitive measures. Here, only one study examined a larger sample (*N* = 18^32^; *N* = 22^34^; *N* = 50^33^). Instead, the model is often used to explain differential effects in individuals without explicitly testing for an inverted U-shaped relatio-ship (*N* = 87^24^; see also^35^).

To bridge this gap, we assessed how enhancing DA neurotransmission via the DA precursor L-DOPA modulates temporal discounting in a large sample. To shed further light on the inconsistent findings reported above, we assessed the influence of individual baseline differences in frequently used putative proxies of DA function, including spontaneous eye blink rate (sEBR)^36^, WM capacity^31^, and impulsivity^24^, on drug effects. SEBR has been linked to DAergic striatal activity^37^, however, the exact neural circuitry through which DA modulates EBR remains open to further investigation. The modulation may involve the spinal trigeminal complex^38^, with DA potentially modulating blinking via basal ganglia pathways, the superior colliculus and nucleus raphe magnus^35^. WM capacity relies on the function of prefrontal neurons that are modulated by the activation of D1 and D2 receptors^39,40^. Impulsive choice may depend on corticostriatal function, i.e. on the balance between cortical and striatal DA tone^20,41^. However, although these measures have often been linked to DA function^,^ more recent PET imaging studies found no evidence for an association between DA synthesis capacity and sEBR, WM capacity, and impulsivity^42,43^.

Our approach expanded upon previous work in several aspects. First, we assessed temporal discounting for both high- and low-magnitude rewards (magnitude effect)^44,45^, to explore the possibility for DA to modulate the magnitude effect, thereby offering deeper insights as opposed to previous studies that focused on either high or low magnitudes. Second, taking advantage of methodological development in cognitive modelling, choices and response times (RT) in our study were simultaneously modelled using a joint temporal discounting drift diffusion model (DDM) – a sequential sampling model for two-alternative forced-choice tasks^18,44^. Based on previous work in humans and animals, we hypothesised DA to reduce rather than increase the rate of temporal discounting (see also preregistration at https://osf.io/a4k9j/). In a second step, we aimed to comprehensively assess individual differences with respect to the DA proxy mea-sures. Our DDM modelling approach then also allowed us to test several additional pre-registered hypotheses regarding effects of DA on response vigour (reduction of RTs and non-decision times)^18,33,46^ and the speed-accuracy trade-off (decision threshold parameter in the DDM)^47^.

## 2 METHODS

### 2.1 Procedure

The study was realised as repeated-measures within-subject design (see also preregistration at https://osf.io/a4k9j/.) After passing a medical assessment to check for contraindications (see section 2.2) by a physician, eligible participants were invited to three separate testing sessions. First, the participants underwent a baseline screening, involving the recording of sEBR (using a webcam, followed by manual rating), testing of WM capacity, including digit span^48^, listening span^49^ and operation span^50^, a comprehensive psychometric assessment (reported elsewhere, see also preregistration at https://osf.io/a4k9j/) including impulsivity (Barratt Impulsiveness Scale, BIS-15^51,52^) and the collection of demographic information.

Following the baseline screening, the participants took part in two double-blind placebo-controlled testing sessions during which they received either L-DOPA or placebo. The measurements were carried out at one-week intervals at a similar time of day and not during the pill break (i.e. the 7-day period in the monthly cycle of hormonal contraceptive use when the user stops taking the contraceptive pill). In individual cases, the interval was shortened or extended due to health or organisational reasons. The participants were instructed to fast for at least 2 h prior to testing. Before each drug session, they were tested for pregnancy, alcohol and drug consumption (THC, cocaine, MDMA, amphetamine, benzodiazepine, barbiturate, methamphetamine, morphine/heroin, methadone and tricyclic antidepressants), and excluded if tested positive. 30 minutes prior to testing (to ensure maximum plasma concentration during the testing phase), the participants received a tablet containing either 150 mg L-DOPA or placebo (maize starch). Physiological parameters and well-being were monitored throughout the entire experimental session. In a follow-up query, the participants could not guess the current drug condition (χ2[1, *N* = 76] = 0.47, *p* = .49).

After the waiting period, the participants performed an incentivised intertemporal choice task, a reinforcement learning task (reported elsewhere) and a visual pattern perception task^53^. The participants were financially reimbursed for participation and received an additional variable bonus depending on task choice. The study was approved by the local ethics committee of the Faculty of Medicine of the University of Cologne.

### 2.2 Sample

We included healthy male and female volunteers aged 25 to 40 who met the following criteria: normal weight as determined by a body mass index (BMI) between 19-25 +/-2 (since the BMI does not account for body composition^43^), righthandedness, normal or corrected-to-normal vision, German as first language (or profound German language skills), and for women hormonal contraception (to prevent pregnancy and reduce hormonal fluctuations). The participants were recruited via university bulletins, mailing lists and by word-of-mouth recommendation.

General contraindications were as follows: intake of common and prescription drugs, participation in other studies involving medication, alcohol or drug intoxication or abuse, strongly impaired vision or strabismus, acute infections, pregnancy, strong emotional burden or physical stress during the study period, past or current psychiatric disorders, neurological disorders, metabolic disorders, internal diseases, chronic pain syndrome, and complications of anaes-thesia (according to self-report). Contraindications specifically regarding the intake of L-DOPA were hypersensitivity to L-DOPA or benserazide, intake of non-selective monoamine oxidase inhibitors, metoclopramide or antihypertensive medication (e.g. reserpine), disorders of the central dopaminergic system, e.g. Parkinsonian Syndromes, increased intraocular pressure (glaucoma), and breastfeeding. We initially aimed for a target sample size of *N* = 100, however, due to COVID-19-related restrictions and challenges, we were unable to reach the final recruitment goal. In total, 85 participants were invited to participate in the study. Nine participants did not complete the study (*n* = 3 due to side effects such as nausea, and *n* = 6 for other reasons such as illness and organisational reasons). This resulted in a final sample of *N* = 76 participants (*n* = 32 female), with a mean age of 28.26 (*SD* = 3.19, range 25 to 40), on average 16.43 years of education (*SD* = 2.30, range 10 to 18), a mean BMI of 22.54 (*SD* = 1.90, range 18 to 26), and average BIS-15 scores of 30.87 (*SD* = 5.03, range 17 to 41). For the assessment of a modulation of the drug effect on temporal discounting by the putative DA proxy measures, *n* = 75 participants were included in the regression analysis, since data for the BIS-15 was missing for one participant due to technical issues.

### 2.3 Temporal discounting task

The temporal discounting task consisted of 192 trials of intertemporal choices, i.e., choices between smaller-but-sooner (SS) and larger-but-later (LL) rewards, interspersed with two short breaks. The task comprised two magnitude conditions (half of the trials each), such that the SS reward was either low (10 €, low-magnitude condition) or high (20 €, high-magnitude condition). The SS reward was always available immediately ("now"), and remained unchanged throughout the experiment). The LL reward consisted of combinations of sixteen multiples of the SS reward value (1^st^ drug session: [1.01 1.02 1.05 1.10 1.15 1.25 1.35 1.45 1.65 1.85 2.05 2.25 2.65 3.05 3.45 3.85], 2^nd^ drug session: [1.011.03 1.08 1.12 1.20 1.30 1.40 1.50 1.60 1.80 2.00 2.20 2.60 3.00 3.40 3.80]) and was available after 6 different delay periods in days (1st drug session: [1 7 13 31 58 122], 2nd drug session: [2 6 15 29 62 118]).

The options were presented in white colour against a black background. An option was selected via button press ("left arrow" or "right arrow" of keyboard) using the right hand, with a RT limit of 4 s. If no button was pressed within this time window, the participant was reminded to decide faster. The chosen option was highlighted with a white frame for 1 s. Trial order and side of option presentation (left/right side of the screen) were randomised. The trials were interspersed with inter-trial intervals of random duration drawn from the interval between 0.5 and 1 s. Beforehand, the participants were informed that upon completion of the task, they would receive the payout from one randomly selected trial, with a delay of the respective choice as a monetary voucher. The task was implemented with MATLAB (The MathWorks, Inc.) and the Psychophysics Toolbox extension^54,55^.

### 2.4 Analysis

#### 2.4.1 Temporal discounting model

We modelled intertemporal choice behaviour using the hyperbolic discounting model^56,57^. To model the choices in the low- and high-magnitude condition within a single model, we fitted a subject-specific parameter *k* reflecting the discounting rate in the low-magnitude condition, and a subject-specific parameter *s*, reflecting the change of the discount rate from the low- to the high-magnitude condition. The subjective (discounted) value (*SV*) of the delayed reward on a given trial *t* was then modelled as follows:

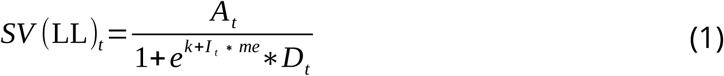

*A* is the amount of the LL reward on trial *t* . Here, *k* is the discount rate (in logarithmic space) for the low-magnitude condition, and *me* models the change in log (*k*) from the low to the high-magnitude condition. *I* serves as an indicator variable for the respective condition (zero for low-magnitude trials, one for high-magnitude trials). *D* specifies the delay of reward receipt.

#### 2.4.2 Softmax choice rule

First, we applied the softmax choice rule^58^ to model the probability of choosing the LL reward on trial *t* as follows:

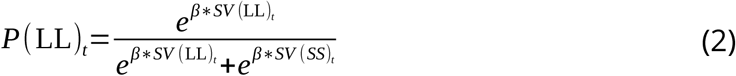

*β* is an inverse temperature parameter, describing the stochasticity of the choices (random choices are described by *β* = 0, while stronger value-dependent choices are described by higher positive *β* values). *SV* is the subjective value of the LL reward on trial *t* . We implemented a single hierarchical Bayesian softmax model (see parameter estimation section below) including the data from both drug conditions, in which the single-subject parameters were drawn from group-level Gaussian distributions. We assigned uniform priors covering numerically plausible values for the group level means and standard deviations of log (*k*), *s* and *β*, and normal priors for parameters modelling changes from the low- to the high-magnitude condition, and changes from the placebo to drug condition.

#### 2.4.3 Drift diffusion model as choice rule

Further, following several recent studies^47,59,60^, we applied the drift diffusion model (DDM) to jointly model the participants’ choices and RTs. For the model’s boundary definitions, the lower boundary was defined as choosing the SS reward and the upper boundary defined as choosing the LL reward. For this purpose, choices towards the lower boundary were multiplied by -1. Since single fast trials may lead to parameter adaption to these trials and hence to a poor model fit at the single-subject level, we excluded the slowest and fastest 2.5% of each participant’s choices from the analysis^61^. The RT on a trial *t* is then modelled using the Wiener first passage time (WFPT):

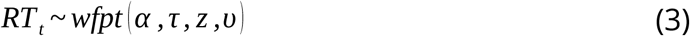

The parameter *α* denotes the boundary separation (speed-accuracy trade-off), *τ* reflects the non-decision time (processing time unrelated to the decision process), *z* reflects the starting-point bias (bias towards one of the two boundaries, with *z* ∈[0,1], i.e. *z*=0.5 reflects a neutral bias and values *z*≥0.5 reflect a bias towards choosing the LL reward), and *υ* denotes the drift rate (rate of evidence accumulation). We fitted three different types of hierarchical DDMs to address different modelling questions, as outlined in the following sections.

##### Condition-specific multivariate DDMs

First, *condition-specific multivariate DDMs* were fitted to the data from each drug condition separately. These models differed in the way in which the trial-wise subjective values modulated the drift rate and were used for model comparison.· For perceptual decision-making, the drift rate varies relative to the strength of evidence^62^. For value-based decision-making, numerous studies have shown that the drift rate is modulated by decision conflict, i.e. the value difference between available options^59,61^. To begin with, we examined a null model (DDM_null_) in which all parameters (*α* , *τ* , *z*, and *υ*) were held constant across trials. Next, to link the hyperbolic discounting model (see equation 1) to the DDM, we implemented two DDMs using different mapping functions linking trial-by-trial variability in the drift rate to trial-wise differences in subjective values as per equation 1. The first of these models was a linear model (DDM_linear_)^63^:

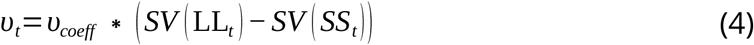

The parameter *υ_coeff_* maps the value differences linearly onto the drift rate *υ* and converts them to the scale of the DDM^63,64^. The second model entailing a value difference modulation was a sigmoid model (DDM_sigmoid_), applying a non-linear transformation of the scaled value differences using an *S*-shaped function^65^:

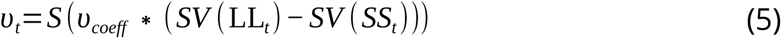

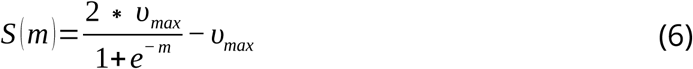

*S* denotes a sigmoid function centred at zero with slope *m* and asymptote *±υ_max_*. All models were implemented as hierarchical Bayesian models, in which the single-subject parameters were drawn from multivariate group-level Gaussian distributions (see parameter estimation section below), with two means corresponding to the magnitude condition (low-magnitude and high-magnitude). The group-level means and standard deviations for the parameters *α* , *τ* , *z*, *υ_max_*, *υ_coeff_* and log (*k*) were assigned uniform priors covering numerically plausible ranges. The parameter *z* (starting-point bias) was estimated in standard normal space, but was backtransformed to the interval [0,1] at the trial level using the *Φ*-transform (standard normal cumulative distribution). These three DDMs (DDM_null_, DDM_linear_ and DDM_sigmoid_) were compared based on approximate leave-one-out (LOO) cross-validation (using Pareto smoothed importance sampling^66^, separately for the two drug conditions.

##### Full multivariate DDM

Second, after confirming that the model ranking (see above) was not affected by the drug condition, we fitted a *full multivariate DDM* including the full covariance matrix of all parameters across all conditions for the best-fitting model. This model was used to obtain posterior estimates of test-retest correlation coefficients for each model parameter, to verify the degree of reliability of each model parameter across drug conditions. Here, each parameter was drawn from a multivariate Gaussian distribution, with four means and four standard deviations for all drug and magnitude conditions (placebo-low, placebo-high, L-DOPA-low, L-DOPA-high) and the full covariance matrix.

##### Drug-effect DDM

Third, we fitted a (univariate) *drug-effect DDM* for the best-fitting model identified using the *condition-specific DDMs* (see above), in which the placebo condition was modelled as the baseline, and for each parameter, drug effects were modelled as additive condition-effect parameters. The purpose of this model was to directly quantify the magnitude of drug-induced changes in each model parameter.

##### Regression model

Lastly to analyse the effect of the putative proxies of DA function (BIS-15 score, sEBR and WM capacity) on the model parameters, we tested for linear and quadratic (inverted U-shaped) modulations of the drug-effect parameters from the full model across drug and magnitude conditions via Bayesian regression. WM capacity was operationalised as principal component (PC) across all three WM tasks (see section S6a of the Supplementary Material). As preregistered for the case of correlated WM measures, we used principal component analysis (PCA) to obtain a compound score reflecting WM capacity. Since BIS-15 scores, sEBR and the first PC of WM capacity were only weakly correlated (-0.08 < *r* < 0.16; see section S6b of the Supplementary Material) we then used each of these proxy measures as individual regressor instead of a compound score. As an example, the regression model is shown below for the drug effect on log ( *k*) as outcome and the putative DA proxies BIS-15 score, sEBR and the first PC of WM capacity as predictors:

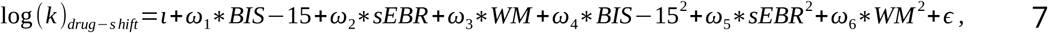

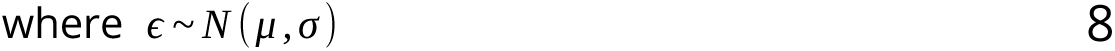

The parameter ι denotes the intercept, _ω1−6_ represent the regression coefficients for the three linear and squared proxy measures BIS-15 score, sEBR and WM capacity, respectively. Note that to avoid double use of parameter names, the conventional naming of the intercept as *α* and regression coefficients as *β*s was changed (*β* denotes the temperature parameter of the softmax model, *α* denotes the boundary separation parameter of the DDM).

##### Analysis of posterior distributions

To quantify the evidence for effects of drug and magnitude on model parameters, as well as effects of the putative DA proxy measures on model parameters, we examined the highest posterior density intervals (HDI) of the posterior distributions of the drug-effect DDM and of the regression coefficients for linear and quadratic terms from the regression model. Here, eEffects were considered reliablecredible if zero fell outside the 95% HDI.

#### 2.4.4 Parameter estimation

Parameter estimation of all models (softmax and DDMs) was performed with Markov chain Monte Carlo (MCMC) simulation, as implemented in Stan^67^ (version 2.26.1), using R^68^ (version 4.3.1) and the R interface to Stan RStan^69^ (version 2.26.23). For the softmax model, sampling was performed with 4,000 iterations for 4 chains (2,000 warmup samples). For the *condition-specific DDM and full multivariate* DDMs, sampling was performed with 8,000 iterations for 4 chains (6,000 warmup samples). For the *drug-effect DDM*, sampling was perfomed with 12,000 iterations for 4 chains (10,000 warmup samples, no thinning). For the regression model, sampling was performed with 2,000 iterations for 4 chains (1,000 warmup samples). The convergence of the chains for the single-subject and group-level parameters was assessed by inspecting the traces such that *R̂* ≤ 1.01^70,71^. For the *condition-specific multivariate DDMs*, as well as for the *drug-effect DDM* we slightly relaxed criteria with regard to *R̂* as compared to the preregistration (for the *drug-effect DDM*, two rather peripheral parameters, the standard deviations of *υ_coeff s_* and *^me^_s_*, had an *R̂* of 1.03 and 1.01, respectively). Inspecting the posteriors confirmed that in all cases, distributions were clearly peaked.

#### 2.4.5 Posterior predictive checks

To ensure that the best-fitting model captures the characteristics of the observed data well, we conducted posterior predictive checks^72,73^. For the softmax model, we ran 4k simulations per trial and overlaid the predicted onto the observed choices. For each of the *condition-specific multivariate DDMs*, we simulated 1k data sets based on the respective posterior distributions. For each participant, we then overlaid the predicted onto the observed RT distributions and LL choice proportions across five LL value bins to capture the associations between choices, RTs and subjective discounted values.

## 3 RESULTS

Descriptive statistics and results from model-agnostic analyses and softmax model are shown in section S1 to S3 of the Supplementary Material, respectively. Numerically, participants made more LL choices under L-DOPA compared to placebo, but this difference was not significant (see section S1 of the Supplementary Material). The hierarchical softmax model revealed a small but reliable effect of L-DOPA on log (*k*), such that log (*k*) was reliably reduced under L-DOPA compared to placebo (see section S2 of the Supplementary Material).

### 3.1 Drift diffusion model comparison

Comparing the *condition-specific multivariate DDMs* with regard to predictive accuracy using LOO cross-validation^66^ confirmed that the data were best accounted for by a DDM including a non-linear, i.e., sigmoid, scaling of the drift rate by the subjective value differences (DDM_sigmoid_). This was the case for both drug conditions (see Table 2). Inspecting the fitted covariance matrix of the *full multivariate DDM* for all parameters and conditions confirmed excellent test-retest reliability for log (*k*) and *υ_coeff s_* , moderate reliability for *υ_max_* and the non-decision time *τ* (see section S5 of the Supplementary Material).

**Table 2.**
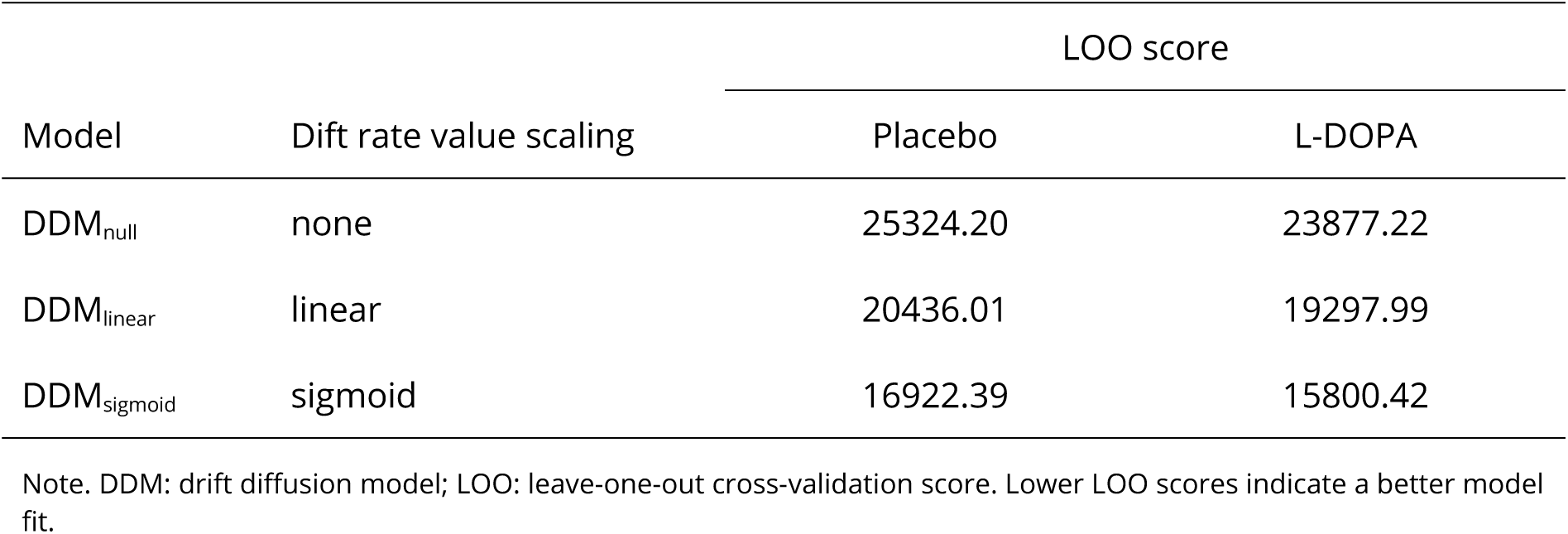
Model comparison for the condition-specific multivariate DDMs.

### 3.2 Posterior predictive checks

Posterior predictive checks for the *condition-specific multivariate DDMs* confirmed that the DDM_sigmoid_ provided a good account of the relationship between RT and LL choice proportions and subjective value (see Figure 1 and 2) in individual participants across the full range of log (*k*) values observed. Further, the DDM_sigmoid_ predicted the choices with the highest accuracy (see Table S4 of the Supplementary Material). In line with the model comparison, this was the case for both drug conditions. Posterior predictive checks for the softmax model are provided in section S3 of the Supplementary Material. Next, we implemented a DDM_sigmoid_ including both magnitude and drug conditions, by including additive drug-effect parameters for capturing changes in parameter values from the low- to high-magnitude and from the placebo to L-DOPA condition. All further analyses are based on the *drug-effect* DDM_sigmoid_.

**Figure 1.**
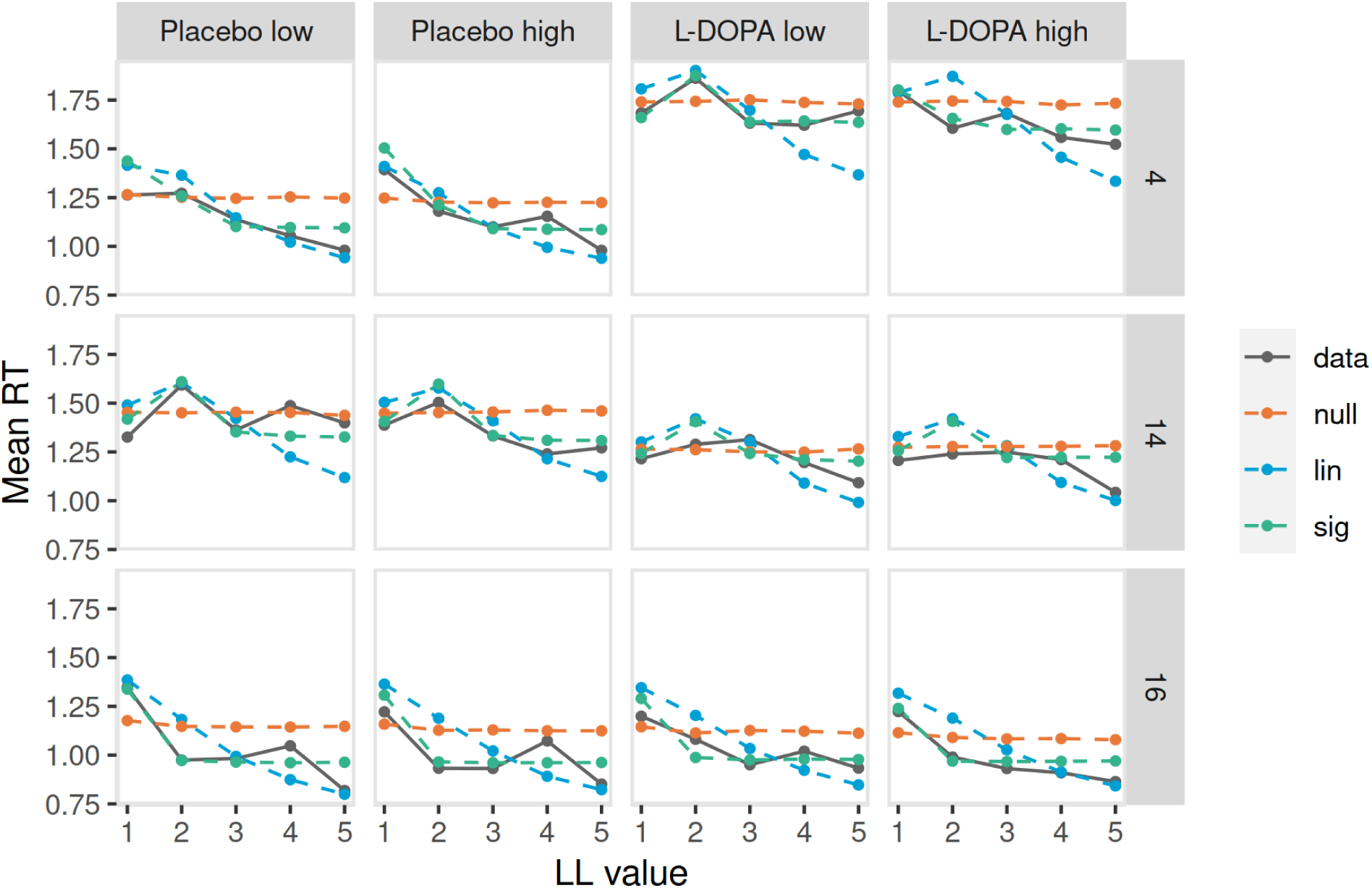
Posterior predictive response time plots for the condition-specific multivariate drift diffusion models (DDM) variants (null: DDM_null_ [orange], lin: DDM_linear_ [blue], sig: DDM_sigmoid_ [green]) for three example participants with a discount rate close to the median (participant #4, top row, *k* = - 5.10), a high discount rate (participant #14, middle row, *k* = -2.64), and a low discount rate (participant #16, bottom row, *k* = -6.77, discount rate from DDM_sigmoid_). Trials were binned into five bins of equal sizes according to the subjective value of the larger-later (LL) option for each participant (based on hyperbolic discounting, see section 2.3.1). The y-axis depicts the observed mean response times per bin (data, grey solid line) and model-predicted mean response times per bin for the different DDMs (coloured dashed lines). Model-predicted response times were generated by averaging over 1k data sets simulated from the posterior distribution of each hierarchical model.

**Figure 2.**
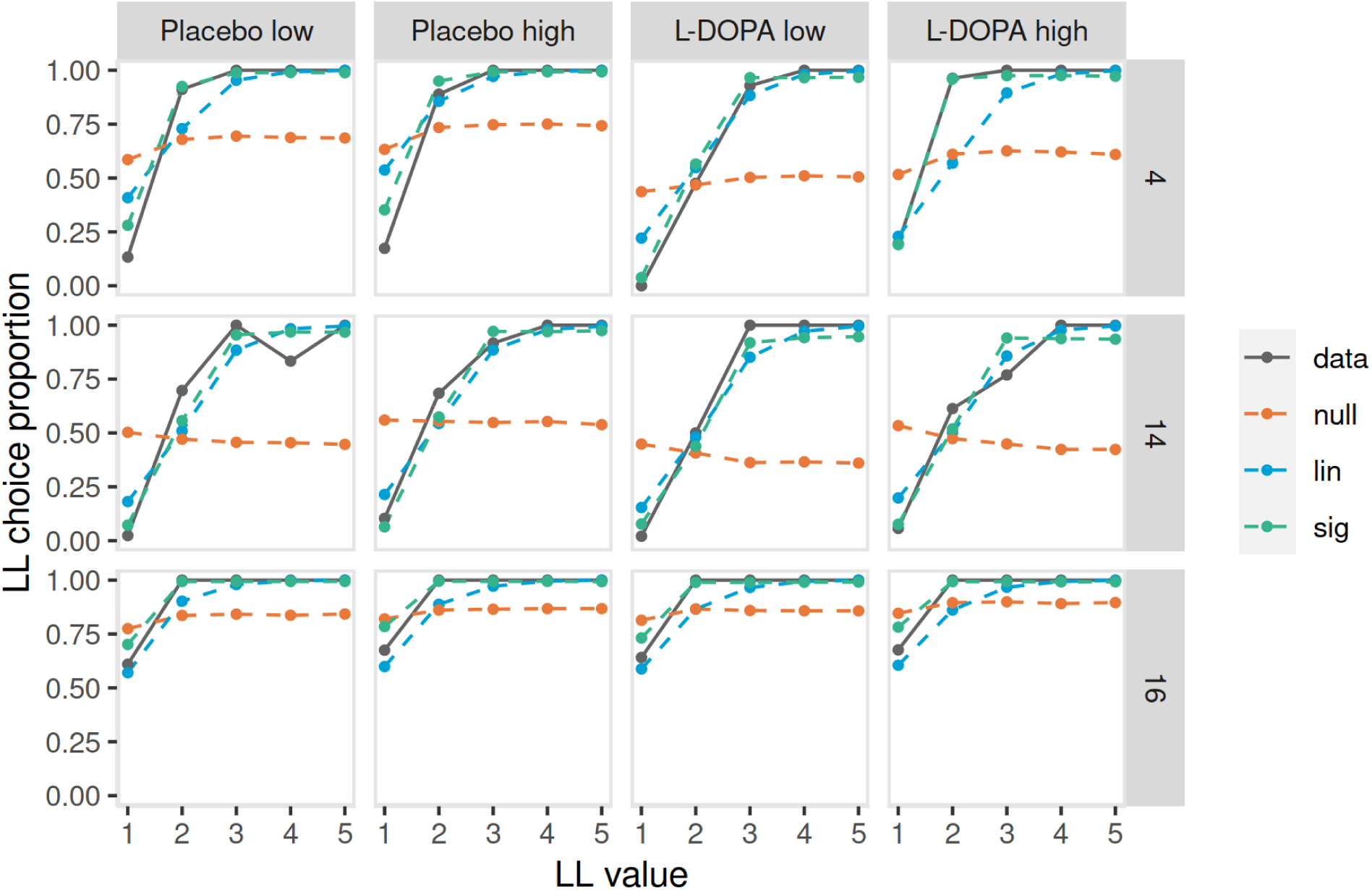
Posterior predictive choice plots for the condition-specific multivariate drift diffusion model (DDM) variants (null: DDM_null_ [orange], lin: DDM_linear_ [blue], sig: DDM_sigmoid_ [green]) for three example participants with a discount rate close to the median (participant #4, top row, *k* = -5.10), a high discount rate (participant #14, middle row, *k* = -2.64), and a low discount rate (participant #16, bottom row, *k* = -6.77, discount rate from DDM_sigmoid_). Trials were binned into five bins of equal sizes according to the subjective value of the larger-later (LL) option for each participant (based on hyperbolic discounting, see section 2.3.1). The y-axis depicts the observed LL choice proportion per bin (data, grey solid line) and model-predicted LL choice proportions per bin for the different DDMs (coloured dashed lines). Model-predicted choices were generated by averaging the proportion of LL choices over 1k data sets simulated from the posterior distribution of each hierarchical model.

### 3.3 Effects of reward magnitude and drug on model parameters

As predicted, higher rewards were discounted less, as indicated by more LL choices in the highcompared to low-magnitude condition (see section S1 of the Supplementary Material for results on model-agnostic analyses), and a negative effect of reward magnitude on log (*k*) (*me*) parameter in the *drug-effect* DDM_sigmoid_, see Table 3 and Figure 3). This magnitude effect on log (*k*) was credibly different from zero (the 95% HDI of the posterior distribution of *me* did not include zero). Likewise, the directional BF_me_^<0^ ^vs. >0^ = Inf provided extreme evidence for values smaller than zero vs. greater than zero.

**Figure 3.**
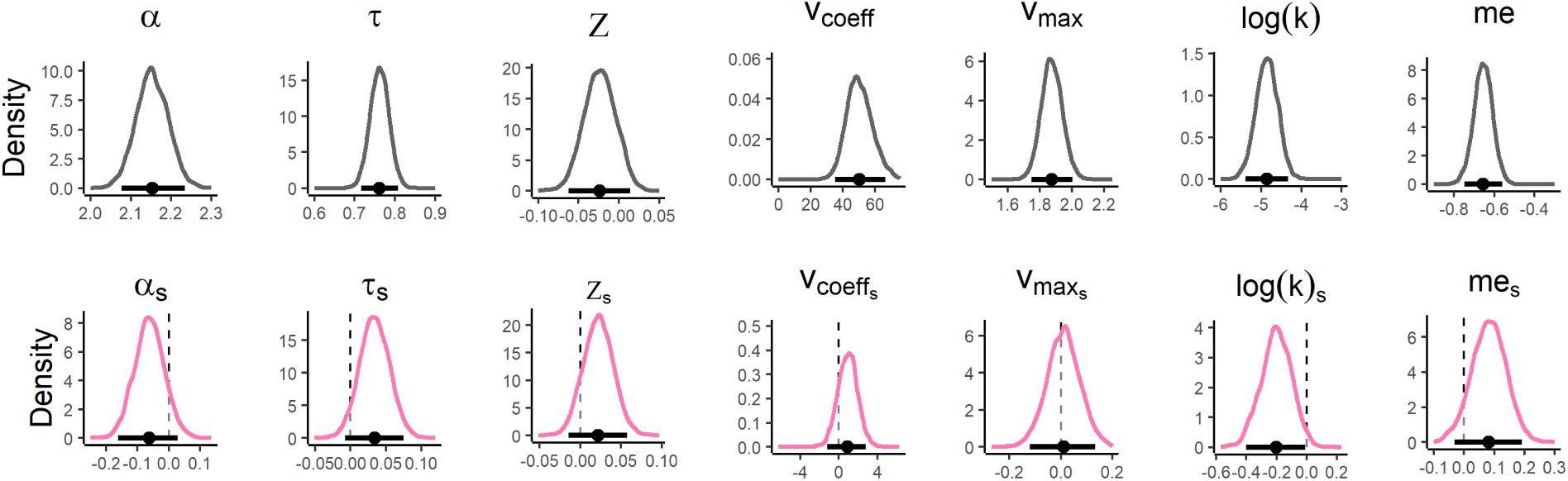
Posterior distributions of the group-level parameter means for the “basic” drift diffusion model parameters and discounting parameters of the drug-effect DDM_sigmoid_. Parameters in the placebo condition (top row, grey distributions) were modelled as baseline, and changes from placebo to L-DOPA (bottom row, pink distributions) as additive shift (s) parameters. The horizontal solid line indicates the 95% highest posterior density interval, the vertical dashed line indicates x = 0. *α* : boundary separation; *τ* : non-decision time; *z*: starting-point bias; *υ_coeff_* : drift rate scaling factor; *υ_max_*: asymptote for *υ_coeff_* , log ( *k*): discount rate; *m e*: magnitude effect for log (*k*); log ( *k*)_s_: drug effect for log (*k*); *me_s_*: drug effect for *m e*.

**Table 3.** Summary of model parameters of the drug-effect DDM (parameter group means and 95% HDIs of the posterior distributions) for the drug-effect DDM_sigmoid_ with parameters in the placebo condition modelled as baseline, and changes from placebo under L-DOPA modelled as additive shift (s) parameters.

|  | Baseline |  |  | Drug effect |  |
| --- | --- | --- | --- | --- | --- |
|  | *M* | 95% HDI |  | *M* | 95% HDI |
| $\alpha$ | 2.15 | [2.08, 2.24] | $\alpha_s$ | -0.06 | [-0.16, 0.03] |
| $\tau$ | 0.76 | [0.72, 0.81] | $\tau_s$ | 0.03 | [-0.01, 0.07] |
| $z$ | -0.02 | [-0.06, 0.01] | $z_s$ | 0.02 | [-0.01, 0.06] |
| $v_{coeff}$ | 50.43 | [35.01, 66.84] | $v_{coeff_s}$ | 0.91 | [-1.17, 2.79] |
| $v_{max}$ | 1.87 | [1.74, 2.00] | $v_{max_s}$ | 0.01 | [-0.12, 0.13] |
| $\log(k)$ | -4.86 | [-5.40, -4.33] | $\log(k)_s$ | -0.20 | [-0.40, -0.01] |
| $me$ | -0.66 | [-0.75, -0.56] | $me_s$ | 0.08 | [-0.03, 0.19] |

Note. HDI: highest posterior density interval; $\alpha$ : boundary separation; $\tau$ : non-decision time; $z$ : starting-point bias; $v_{coeff}$ : drift rate scaling factor; $v_{max}$ : asymptote for $v_{coeff}$ ; $\log(k)$ : discount rate; $me$ : magnitude effect for $\log(k)$ ; $\log(k)_s$ : drug effect for $\log(k)$ ; $me_s$ : drug effect for $me$ .

The posterior distributions of the *drug-effect* DDM_sigmoid_, in which the placebo condition was modelled as baseline and drug effects as additive changes from the baseline, revealed that enhancing DA transmission reduced discounting, as indicated by a negative drug effect on log (*k*) (log (*k*)*_s_*) parameter in the DDM_-sigmoid_, see Table 3 and Figure 3), substantiated by the observation that the 95% HDI of the posterior distribution did not include zero, and BF_log(k)s_^<0vs.>0^ = 49.15 provided substantial evidence for values smaller than zero vs. greater than zero. Recall that these effects were reproduced using the softmax choice rule (see Table S2 and Figure S2 of the Supplementary Material).

In contrast, the drug effect on the magnitude effect (*^me^_s_* parameter) was not credibly different from zero in both the softmax model and DDM (the 95% HDI of the posterior distribution included zero in both cases). For the remaining DDM parameters (*α* , *z*, *τ* , and *υ_max_*), we found no credible evidence for drug effects, since in all cases, the 95% HDI of the posterior distributions included zero (BF*_α_*_s_^<0 vs. >0^ = 8.83; BF*_τ_*_s_^>0 vs. <0^ = 17.35; BF_zs_^>0 vs. <0^ = 7.06; BF*_υ_*_coeff-s_^>0 vs. <0^ = 4.45; BF*_υ_*_coeff-max_^>0 vs. <0^ = 1.28). This suggests that these parameters were not credibly modulated by the drug manipulation (see Table 3, column 2 and Figure 3, bottom row).

### 3.4 Effects of interindividual differences

To test for a modulation of the drug effects by interindividual measures of putative DA proxies, we regressed sEBR, the first principal component of a PCA across the working memory (WM) tasks (digit span^48^, listening span^74^, and operation span^75^) and impulsivity (BIS-15 score^76,77^) onto the drug-effect parameters of the *drug-effect* DDM_sigmoid_. We found no credible evidence that any of the drug effects were substantially modulated by the DA proxy measures, neither in a quadratic (see Figure 4 and Figure S6d of the Supplementary Material) nor linear manner (see Figures S6d and S6e of the Supplementary Material), since for all posterior distributions of the regression coefficients, the 95% HDIs included zero (see Table S6e for the BFs for directional effects).

**Figure 4.**
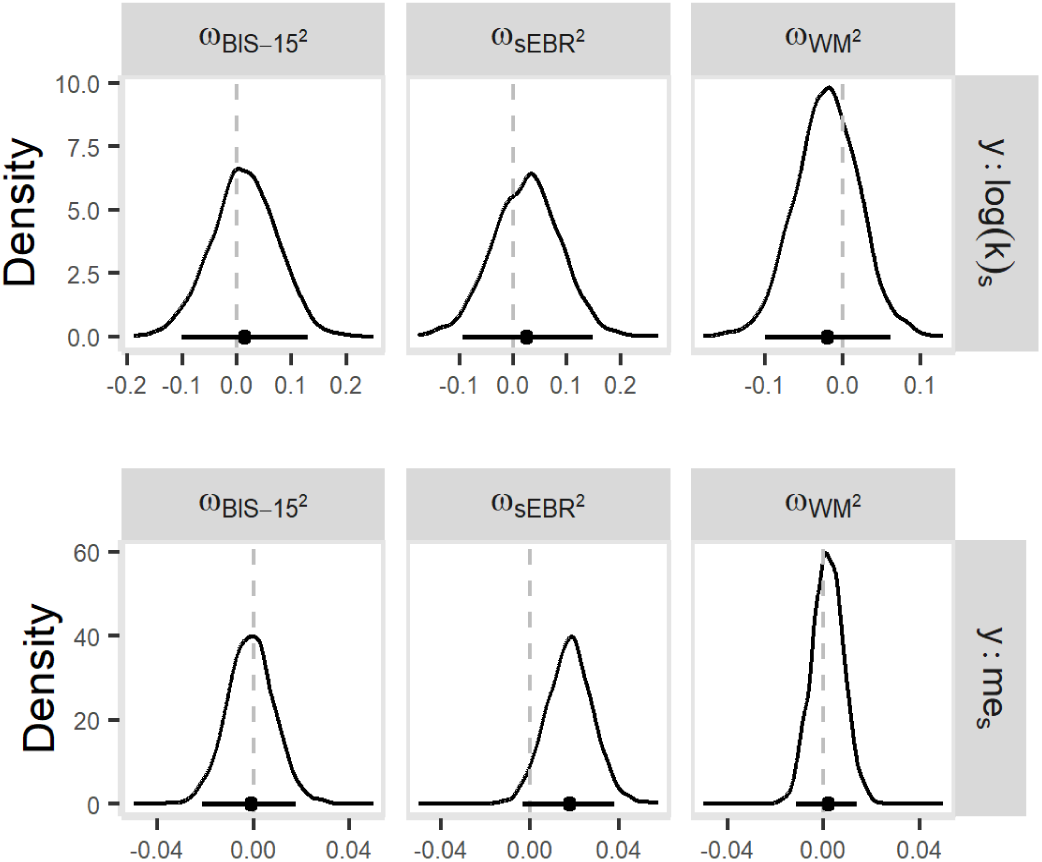
Posterior distributions of the quadratic coefficients (_℧_) for the dopamine proxies impulsivity (BIS-15 scores), spontaneous eye blink rate (sEBR) and working memory capacity (WM), regressing the proxies onto the drug-shift (s) parameters for the discount rate log ( *k*) and the magnitude effect *m e* of the drug-effect DDM_sigmoid_. The posterior distribution of the intercept is not depicted. The horizontal solid line indicates the 95% highest posterior density interval, the vertical dashed line indicates x = 0. log ( *k*)_s_: drug effect for log (*k*); *me_s_*: drug effect for *m e*.

## 4 DISCUSSION

In the current work, we revisited the role of dopamine (DA) neurotransmission in temporal discounting, and the role of interindividual differences in putative proxies for DA function, including impulsivity, WM capacity and sEBR. Previous pharmacological studies in humans have yielded highly heterogeneous results, which may be attributable to meaningful interindividual differences in baseline DA function, and/or noisy effects due to small samples. Here we applied comprehensive computational modelling using temporal discounting drift diffusion models (DDM) to dissociate effects of value magnitude, DA and interindividual differences on decision-making.

As predicted and replicating previous findings^59,78^, we found a magnitude effect for temporal discounting, such that higher rewards were discounted less, supported by both the model-agnostic analysis and computational modelling. We found no credible drug effect on the magnitude effect – if anything, L-DOPA slightly dampened the effect (but the 95% HDI overlapped with zero).

Likewise a recent study entailing a re-analysis of pharmacological intervention data examined effects of DA D1 receptor stimulation via the DA agonist PF-06412562 and DA D2 receptor blockage using amisulpride on DDM parameters^79^. While DA D1 stimulation and DA D2 receptor blockade had no effect on temporal discounting (discount rate k), DA D2 blockade impacted on the drift rate, increasing sensitivity to reward magnitude during evidence accumulation. Further, while the starting point was closer to the LL boundary under placebo, this bias was reduced by DA D2 blockade. Observing no effect of D1 agonism on temporal discounting stands in contrast to our findings. However, the results may not be directly comparable to ours, since substances with different receptor affinities were used (PF-06412562 being a selective D1 agonist and L-DOPA interacting with both D1 and D2 receptors^80,81^). Further, we found no effects of L-DOPA on drift rate value scaling or starting point. Nevertheless, although the authors did not observe any effect on the discount rate following decreased DA neurotransmission through D2 blockade, they did find an impact on the drift rate, such that D2 blockade increased evidence accumulation for LL choices for large differences in reward magnitude between the LL and SS options. This finding bears some resemblance to our own results, though in an inverse manner, as we observed a reduction in the discount rate when increasing DA neurotransmission using L-DOPA, indicating a reduced tendency to devalue LL rewards. Further, as observed in our study, drug effects did not relate to WM capacity^79^.

Notably, no drug effects on model parameters were reliably modulated by individual differences in putative DA proxy measures. We observed a reliable reduction in discounting following administration of L-DOPA. This effect was small (i.e., about one-third of the size of the magnitude effect) but nonetheless challenges results from influential previous studies, for example Pine and colleagues^82^, reporting increased temporal discounting under L-DOPA in a sample of *N* = 13 participants. Furthermore, it has been proposed that impulsive choice (i.e., steeper temporal discounting) is linked to higher dopaminergic tone^83,84^. However, the evidence from human studies is rather mixed. While some studies report an increase in impulsive choice when enhancing DA transmission^82^, some report a decrease in impulsive choice^20,29,60^ or no overall effects^22,24,85^. Our results are therefore more in line with findings from animal studies^86,87^ as well as several more recent human studies^18,19,88^ suggesting that DA reduces, rather than increases, temporal discounting, albeit with a small effect size. DA is believed to impact on the cost-benefit trade-off by increasing tolerance to costs and/or facilitating overcoming costs by increasing vigour when maximising overall subjective re-wards^89–91^. From this it would follow that DA would reduce the effect of waiting cost, thereby increasing the preference for LL over SS rewards in intertemporal choice, which fits with our results of reduced discounting under L-DOPA compared to placebo.

Inconsistencies with other findings may in part be attributable to low sample sizes in earlier studies (*N* = 22^85^, *N* = 10^22^; *N* = 23^20^; and *N* = 13^82^), or effects being modulated by meaningful interindividual differences in cognitive functions purportedly related to DA levels, e.g. impulsivity^92^. Data from animal and human studies suggest that differential effects of DA administration may reflect variability in cognitive functions related to baseline DA levels^35,93^, leading to the highly-cited assumption that the relationship between WM capacity and DA level follows an inverted U-shaped function, such that depending on baseline DA levels, dopaminergic drugs may elicit opposing effect (for a review, see ^94^). However, we did not observe credible evidence for associations of the interindividual difference measures with DA drug effects, neither for sEBR nor WM capacity or self-reported impulsivity, contrasting with some previous results in smaller samples. Several reasons may explain the absence of such a relationship. First, an inverted U-shaped function may not be the appropriate model for the relationship between the individual difference measures and drug effects – we focused on linear and quadratic effects but did not explore other functions^94^. Second, some of the studies supporting an inverted U-shaped model had comparatively small sample sizes (*N* = 18^32^ and *N* = 22^34^) or explained heterogeneous findings for subgroups in regard of a quadratic relationship without explicitly testing for this^92^.

Consistent with the absence of an inverted-U-function, two studies measuring D2 receptor availability with PET imaging found no evidence for a relationship with sEBR (*N* = 20^95^ and *N* = 30^96^). Also, a recent PET imaging study in a large sample of healthy participants (*N* = 94^97^) found no evidence for the proposed U-shaped relationship between sEBR, WM capacity and DA D2 receptor availability.

This suggests that heterogeneous earlier findings may have resulted from noisy estimates obtained in underpowered studies. Nonetheless, other measures of interindividual differences may be better proxies of DA function, or the relationship may be mediated by DA release or DA D2/3-receptor availability instead^97^. However, although we did not measure DA function directly, the absence of a modulating DA proxy effect on L-DOPA drug effects in the present sample of *N* = 76 is inconsistent with the idea that these measures directly relate to DA function. Further, if these measures reflect baseline DA function, one would expect them to be correlated. However, this was not the case. Therefore, while these measures may reliably reflect interindividual differences, our results suggest that they do not explain variance in individual differences in DA drug effects on temporal discounting. Similarly, a study assessing the effects of DA on risk aversion found that a D1 agonist increased risk aversion, however, the effect was not modulated by WM capacity^23^. Disparate findings may further be related to the studies differing with regard to the substance employed (buproprion^85^, d-amphetamine^29^, pramipexol^22^, tolcapone^20^, L-DOPA^82,92^, and haloperidol^60^), thereby exerting different mechanisms at the receptor level. For instance, haloperidol, a DA receptor antagonist, which, at lower doses, is assumed to increase striatal DA transmission, reduced delay discounting in two other recent studies^60,98^.

The reduction of temporal discounting under L-DOPA is in line with findings from patients with Parkinson’s disease (PD) exhibiting reduced temporal discounting ON vs. OFF dopaminergic medication^99^. Likewise, DA increases have been linked to improvements in model-based control in healthy volunteers ^100^(but see ^101^) and PD patients^102^. DA might therefore improve cognitive functions supporting both model-based control and temporal discounting, such as planning ^103^ and executive control^104^. Striatal DA may also contribute to decision-making by regulating the activation balance of striatal *go* and *no-go* pathways^105–107^. Increased striatal DA levels may therefore also increase the signal-to-noise ratio of striatal value representations^18^, thereby increasing the likelihood that larger-later rewards gain access to prefrontal cortical processing.

With regard to the overall effects of L-DOPA on the remaining model parameters, we further expected L-DOPA to increase response vigour (i.e., reduce RTs), which should be reflected in shorter non-decision times^60^, and to reduce the decision threshold, reflected in a reduced boundary separation parameter^47,108^. Model-agnostic analysis revealed no significant difference in overall RTs between L-DOPA and placebo. For the boundary separation, the directionality of the effect was in the predicted direction. If anything, L-DOPA reduced the boundary separation, but the effect was small, and the 95% HDI showed some overlap with zero. The estimate of non-decision time (at baseline, i.e. under placebo) was comparable to that of our previous studies^60,61^, suggesting consistency in this parameter across different samples and experimental contexts. Under L-DOPA, the non-decision time was, if anything, slightly increased (although here, the 95% HDI showed overlap with zero). This is inconsistent with earlier results using different dopaminergic drugs^60^. Our findings add to a growing body of evidence that enhancing DA transmission decreases^18,19,88^ rather than increases^82^ impulsive choice as measured by temporal discounting tasks. Further, we found no evidence that interindividual differences in sEBR, WM capacity and impulsivity modulated the drug effects in an inverted U-shaped manner, questioning their frequently proposed role as proxies for baseline DA function^23,71,72,81,82^.

Posterior predictive checks confirmed that the DDM_sigmoid_ accounted best for associations between RT and LL choice proportions and subjective value. In contrast, since the DDM_null_ predicted constant RTs across trials, it failed to reproduce the observed reductions in RT for increasing subjective value. Both the DDM_linear_ and DDM_sigmoid_ reproduced the subjective-value-related reductions in RTs. However, the DDM_linear_ tended to over and underestimate RTs for the extremer ends of LL values, i.e., small and large LL values, respectively. With regard to LL choice proportions, again, the DDM_null_ predicted nearly constant proportions across trials, thereby failing to reflect the increase in LL choices with increasing subjective LL value. Both the DDM_linear_ and DDM_sigmoid_ reproduce this pattern, however the model with the linear mapping more frequently underestimated LL choice proportions. Comparing the fit of the linear and sigmoid DDMs, it is important to note the varying performance of these models across different regions of the data. The sigmoid mapping, in essence, bounds the drift rate, preventing it from becoming excessively large for high value differences. When considering value differences between the LL and SS rewards, both the linear and sigmoid DDM generate similar predictions for the intermediate range of data, whereas the linear DDM will under/overpredict RTs and LL choice proportions for more extreme value differences. This can be seen in the posterior predictive checks for participant #4 and #14 in Figure 1 and participant #4 and #16 in Figure 2. Overall, the DDM_sigmoid_ predicted the choices with the highest accuracy for both drug conditions, and posterior predictive checks demonstrated that this model accounts for the full range of observed data at the individual-participant level. Given the variability observed across participants, these differences in model fit would be obscured by focusing on in group-level predictions.

### Limitations

The current study also has several limitations which need to be addressed. First, we did not assess effects of hormonal cycle on the effect of L-DOPA on temporal discounting. However, given that ovarian hormones interact with DA signalling^109^, and in the light of cycle-dependent fluctuations in decision-making^110^, this factor should be addressed in future studies. Furthermore, the rewards were hypothetical. However, since choice preferences for real and hypothetical rewards and discounting measures derived therefrom show close correspondence. Furthermore, it is also conceivable that the hypothesised moderating effect of the putative DA proxies on the effects of L-DOPA on temporal discounting did not mani-fest because the variance in the measures was rather small. Since we collected data from a healthy control sample, we may not have tapped data points from the extreme ends, e.g. from high impulsive individuals or individuals with low working memory capacity.

### Relevance

It is well established that dopamine is involved in the valuation of future rewards and also regulates various other aspects of decision-making such as the regulation of exploration and exploitation behaviour^60,108,111,112^. Understanding the relationship between dopaminergic signalling and specifically intertemporal is choice vital, since increased devaluation of future rewards is often found in psychiatric disorders characterised by alterations in dopaminergic neurotransmission^6,113^, such as substance use disorders and behavioural addictions^58,114^. Further, dopamine replacement therapy in Parkinson’s Disease patients can lead to impulse control disorders (for reviews, see^115,116^). Therefore, assessing the effects of L-DOPA on decision-making in healthy volunteers is of considerable relevance to delineate underlying mechanisms.

## Conclusions

In the current study, we investigated the effects of increasing DA neurotransmission using the DA precursor L-DOPA on temporal discounting, and the role of interindividual differences in putative proxies of DA function, including sEBR, working memory capacity and impulsivity. Using a model-based approach via temporal discounting drift diffusion modelling, we confirmed that the choices and RTs were best explained by a non-linear, sigmoid scaling of the drift rate by subjective value differences. Contrasting with some earlier findings in substantially smaller samples, L-DOPA reliably reduced temporal discounting across different modelling schemes. Inconsistent with the prominent inverted-U-model, interindividual differences in sEBR, working memory capacity and impulsivity did not ex-plain variance in DA drug effects on temporal discounting, neither in a linear nor inverted U-shaped manner. Our findings therefore suggest that these measures may not be related to baseline DA function, which is further corroborated by the absence of strong intercorrelations. In conclusion, increasing DA neurotransmission reduced temporal discounting, but our findings challenge the utility of using sEBR, working memory capacity and impulsivity as proxies of DA function.

## CODE AND DATA AVAILABILITY

All model code is openly available on the Open Science Framework at https://osf.io/5fkjy/. The fitted models are openly available at https://zenodo.org/records/13378364 (DOI: 10.5281/zenodo.13378364). The data cannot be shared publicly, since the participants did not provide consent to public data access. The data are available from RADAR at https://www.radar-service.eu/radar/en/dataset/rQLffRDOgzeCXGqI (DOI: 10.57743/rQLffRDOgzeCXGqI), whereby access is granted to researchers for scientific purposes only.

## AUTHOR CONTRIBUTIONS

Elke Smith: Conceptualisation, methodology, software, formal analysis, investigation, data curation, visualisation, project administration, writing - original draft, writing - review & editing Hendrik Theis: Resources, investigation, writing - review & editing Thilo Van Eimeren: Conceptualisation, resources, investigation, writing - review & editing Kilian Knauth: Investigation, data curation, writing - review & editing Deniz Tuzsus: Investigation, writing - review & editing Lei Zhang: Methodology, writing - review & editing David Mathar: Methodology, writing - review & editing Jan Peters: Conceptualisation, methodology, project administration, supervision, funding acquisition, writing - review & editing

## ACKNOWLEDGEMENTS

The authors thank Lea Kemalides, Hannah Hacker and Emily Burlon for helping with study organisation and data collection.

