## Supplement for "Dopamine and temporal discounting: revisiting pharmacology and individual differences"

### S1 Model-agnostic analyses

Numerically, the response times and proportion of larger-later (LL) choices were higher for the high- compared to low-magnitude condition, and under L-DOPA compared to placebo (see Table S1 and Figure S1). Response times were not significantly different between the high- compared to the low-magnitude condition ($F(1) = 0.07$, $p = .797$, $\eta_p^2 = 2.22\times10^{-4}$), or between the drug compared to the placebo condition ($F(1) = 0.04$, $p = .844$, $\eta_p^2 = 1.29\times10^{-4}$), and no interaction between the two factors ($F(1) = 0.02$, $p = .878$, $\eta_p^2 = 7.91\times10^{-4}$). The participants made significantly more LL choices in the high compared to the low-magnitude condition ($F(1) = 7.91$, $p = .005$, $\eta_p^2 = 0.03$), but not between the L-DOPA compared to the placebo condition ($F(1) = 0.39$, $p = .535$, $\eta_p^2 = 1.29\times10^{-3}$), with no interaction between the two factors ($F(1) = 0.02$, $p = .885$, $\eta_p^2 = 6.99\times10^{-5}$).

Table S1. Descriptive statistics.

| | Magnitude condition | | | | Drug condition | | | |
|---|---|---|---|---|---|---|---|---|
| | Low | | High | | Placebo | | L-DOPA | |
| | *MD* | *SD* | *MD* | *SD* | *MD* | *SD* | *MD* | *SD* |
| RTs | 1.37 | 0.59 | 1.26 | 0.54 | 1.35 | 0.59 | 1.37 | 0.58 |
| LL choices | 0.61 | 0.22 | 0.71 | 0.20 | 0.66 | 0.22 | 0.69 | 0.21 |

Note. MD: median; RTs: response times; LL: larger-later.

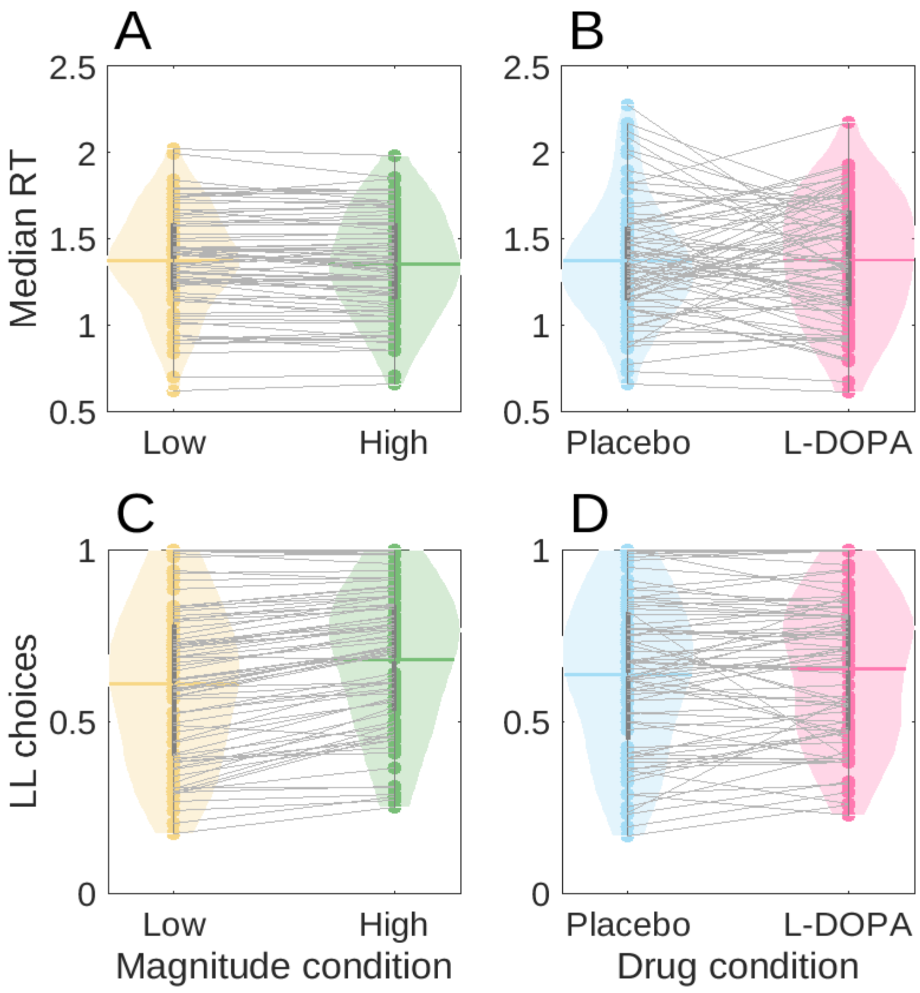


Figure S1. Median response times (RT) per magnitude (panel A) and drug condition (panel B), and proportion of larger-later (LL) choices per magnitude (panel C) and drug condition (panel D). Horizontal thick lines: medians; points and grey lines: individual participant data.

### S2 Softmax choice rule

Modelling the choices with the softmax choice rule, we found reduced discounting for higher rewards (magnitude effect), as indicated by the negative effect of value magnitude on $\log(k)$ ($me$ parameter, the 95% HDI of the posterior distribution not including zero), and reduced discounting under L-DOPA compared to placebo, as indicated by a negative drug effect on $\log(k)$ ($\log(k)_s$ parameter, see Table S2 and Figure S2, the 95% HDI of the posterior distribution not including zero). The drug effect on the magnitude effect was not reliably different from zero (the 95% HDI of the posterior distribution including zero).

Table S2. Group-level means and 95% HDIs of the posterior distributions from the softmax model.

| | *MD* | *95% HDI* |
|---|---|---|
| $\beta$ | 1.75 | [1.26, 2.31] |
| $\beta_s$ | 0.05 | [-0.29, 0.38] |
| log(k) | -4.76 | [-5.31, -4.24] |
| $\log(k)_s$ | -0.24 | [-0.48, -0.02] |
| me | -0.59 | [-0.69, -0.49] |
| $me_s$ | 0.02 | [-0.13, 0.15] |

Note. HDI: highest posterior density interval; β: inverse temperature parameter; βs: drug effect for β; log(k): discount rate, $\log(k)_s$: drug effect for log(k); me: magnitude effect; $me_s$: drug effect for magnitude effect.

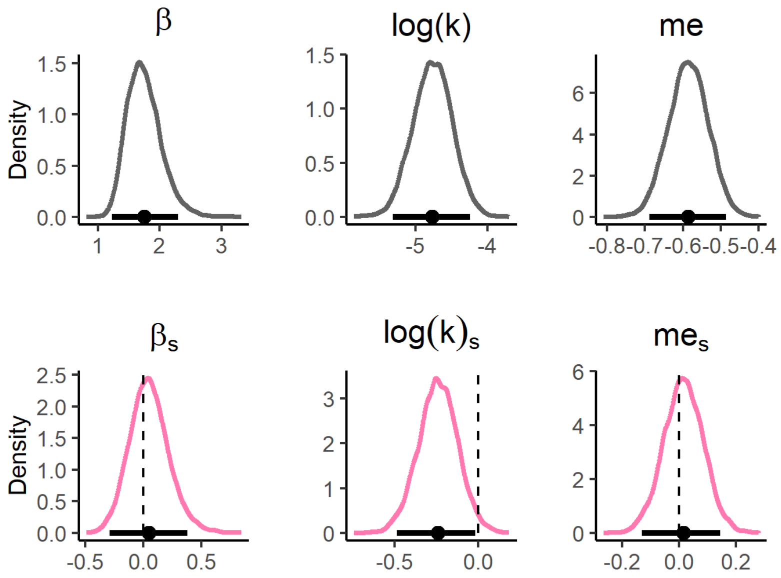


Figure S3. Posterior distributions of the group-level parameter means of the softmax model. The horizontal solid line indicates the 95% highest posterior density interval, the vertical dashed line indicates x = 0. Parameters in the placebo condition (top row, grey distributions) were modelled as baseline, and changes from placebo to L-DOPA (bottom row, pink distributions) as additive shift (s) parameters. $\beta$: inverse temperature parameter; $\beta_s$: drug effect for $\beta$; log(k): discount rate; $\log(k)_s$: drug effect for $\log(k)$; $me$: magnitude effect; $me_s$: drug effect for magnitude effect.

### S3 Posterior predictive checks for the softmax model

To verify that the softmax model can account for the observed choices, we conducted posterior predictive checks. We simulated 4k data sets based on the posterior distributions of the hierarchical softmax model and overlaid the predicted onto the observed choices. This demonstrated that the softmax model accounted well for the observed LL choice proportions (see Figure S3).

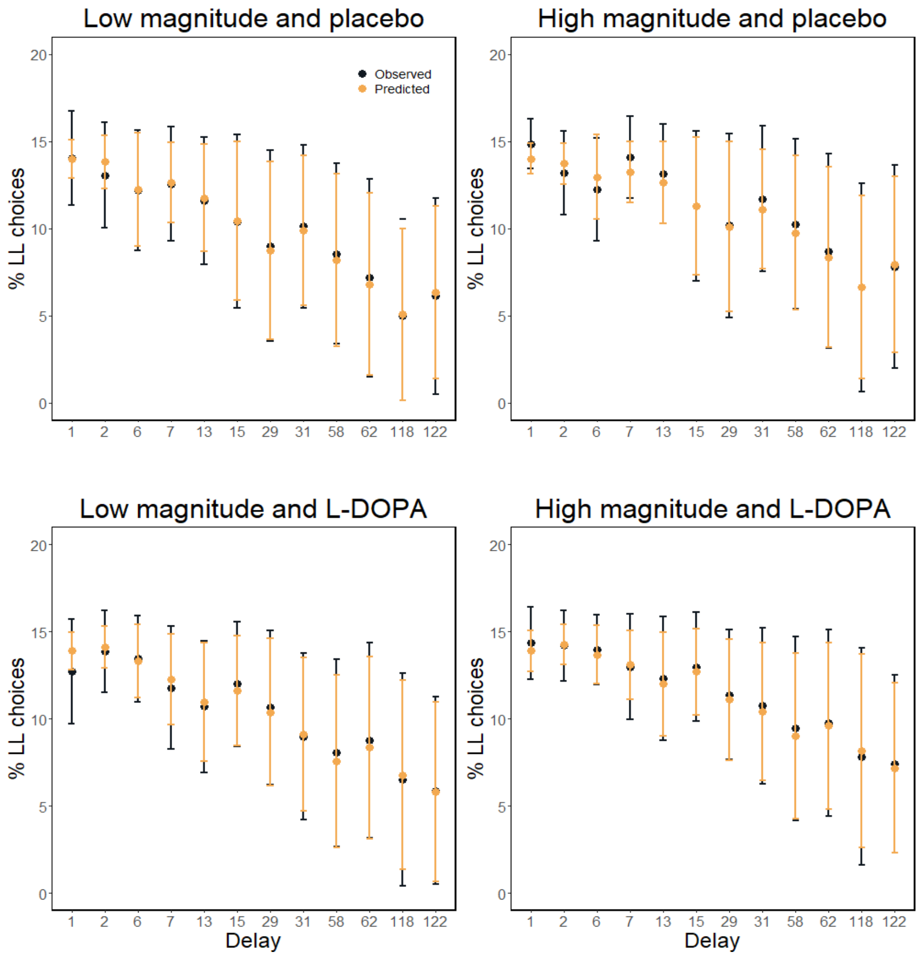


Figure S1. Observed (teal) and predicted (orange) proportions of larger-later (LL) choices across participants split by magnitude and drug condition. Model-predicted choices were obtained by averaging over 4k simulations per trial from the posterior distribution of the softmax model. The scaling of the x-axis has been changed to “factor” for better readability. Points: means; bars: standard deviations.

### S4 Posterior predictive checks for the condition-specific drift diffusion models

Table S4. Accuracy of the choice predictions for the condition-specific multivariate DDMs.

| | Placebo | | L-DOPA | |
|---|---|---|---|---|
| | Low | High | Low | High |
| $DDM_{null}$ | 0.63 | 0.65 | 0.63 | 0.65 |
| $DDM_{linear}$ | 0.74 | 0.75 | 0.73 | 0.75 |
| $DDM_{sigmoid}$ | 0.83 | 0.84 | 0.82 | 0.84 |

Note. Model-predicted choices were generated by averaging the proportion of LL choices over 1k data sets simulated from the posterior distribution of each hierarchical model. DDM: drift diffusion model.

### S5 Reliability of drift diffusion parameter estimates

To estimate the test-retest reliability, we implemented the best-fitting drift diffusion model (DDM), i.e. the $DDM_{sigmoid}$, as a model spanning both drug sessions (*full multivariate DDM*). The model parameters of each individual were drawn from group-level multivariate Gaussian distributions, with means and covariance matrices for all parameters, setting uniform priors from -1 to 1 for the test-retest correlation coefficient parameters (see section 2.3.4 of the main text for details on parameter estimation). Test-retest reliability was excellent for $\upsilon_{coeff}$ (drift-rate scaling parameter) and $\log(k)$ (discount rate) for both magnitude conditions (see Table S5 and Figure S5). For $\alpha$, test-retest reliability was poor for both magnitude conditions. For $\tau$ and $\upsilon_{max}$, test-retest reliability was moderate. Further, while test-retest reliability was good for $z$ (starting-point bias) for the low-magnitude condition, it was poor for the high-magnitude condition. This suggests that the magnitude effect (see section 3.2 of the main text) is mediated by the starting-point bias only in some individuals. While in the low-magnitude condition, the starting point appeared to be consistently biased towards smaller-sooner (SS) rewards, this was not the case in the high-magnitude condition.

Table S5. Means and 95% HDIs of the posterior distributions of the correlation coefficients from the full multivariate DDM between placebo – low magnitude and L-DOPA – low magnitude, and placebo – high magnitude and L-DOPA – high magnitude, respectively.

| | Low magnitude | | High magnitude | |
|---|---|---|---|---|
| | *M* | *95% HDI* | *M* | *95% HDI* |
| $\alpha$ | 0.28 | [0.05, 0.52] | 0.27 | [0.02, 0.51] |
| $\tau$ | 0.55 | [0.39, 0.70] | 0.56 | [0.40, 0.71] |
| $z$ | 0.76 | [0.60, 0.90] | 0.24 | [-0.04, 0.53] |
| $\upsilon_{coeff}$ | 0.98 | [0.95, 1.00] | 0.97 | [0.93, 1.00] |
| $\upsilon_{max}$ | 0.47 | [0.25, 0.68] | 0.60 | [0.42, 0.78] |
| $\log(k)$ | 0.93 | [0.89, 0.96] | 0.93 | [0.88, 0.96] |

Note. DDM: drift diffusion model; HDI: highest posterior density interval; α: boundary separation; $z$: starting-point bias; τ: non-decision time; $\upsilon_{coeff}$: coefficient mapping the value differences onto the drift rate using a sigmoid function $S$; $\upsilon_{max}$: asymptote for $S$; log(k): discount rate.

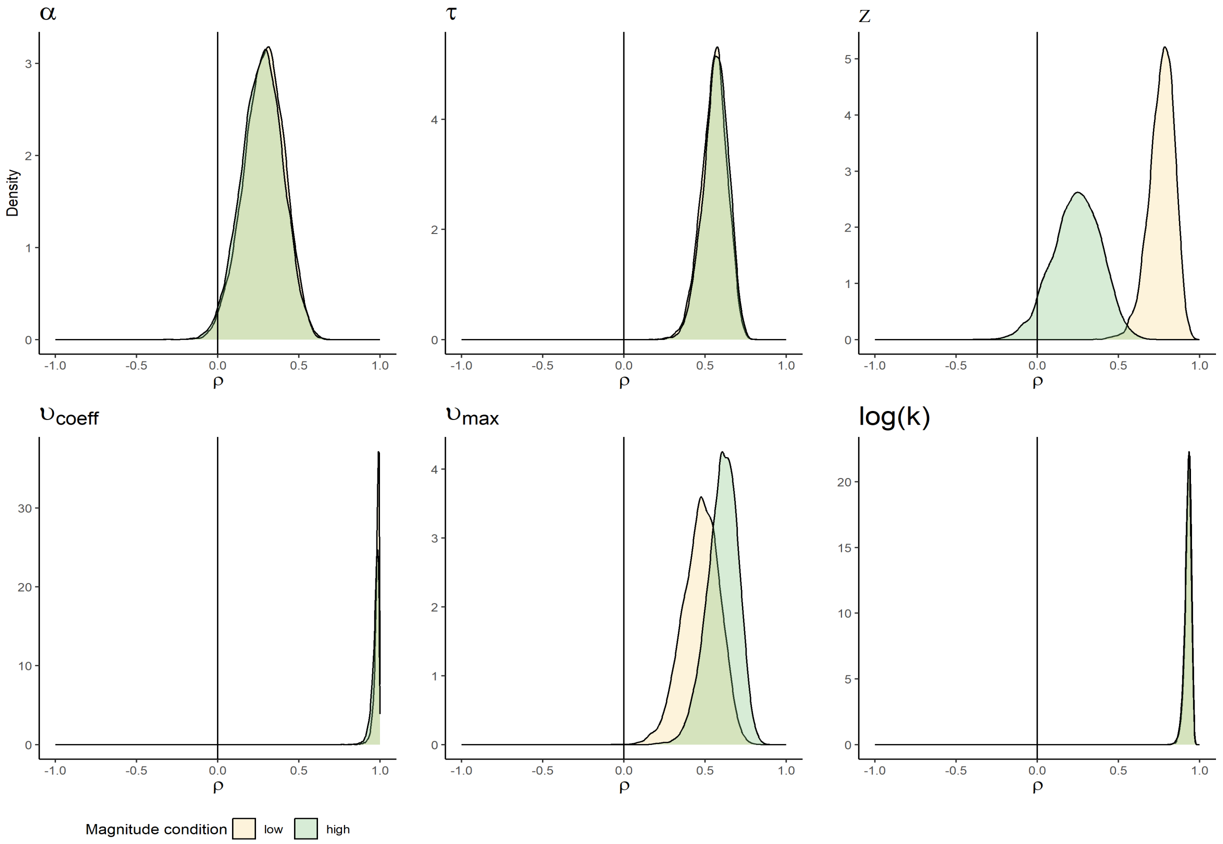


Figure S5. Test-retest (placebo vs. L-DOPA) reliability of the parameter estimates from the full multivariate drift diffusion model (DDM). Posterior distributions of the correlations between placebo – low magnitude and L-DOPA – low magnitude (yellow), and placebo – high magnitude and L-DOPA – high magnitude (green), respectively. $\alpha$: boundary separation; $\tau$: non-decision time; $z$: starting-point bias; $\upsilon_{coeff}$: coefficient mapping the value differences onto the drift rate using a sigmoid function $S$; $\upsilon_{max}$: asymptote for $S$; log(k): discount rate.

### S6 Effects of interindividual differences

#### S6a Principal component analysis for working memory capacity

Working memory capacity was operationalised as principal component across three working memory tasks (digit span [backward and forward], listening span, and operation span). Principal component analysis was performed using MATLAB's (The MathWorks, Inc.) pca function. All scores were *z*-scored before entering the analysis. See Table 6a for the correlations between the working memory scores. The cumulative variance explained and factor loadings are depicted in Figure S6a.

Table S6a. Correlations between the working memory scores.

| | Digit span fw max | Listening span | Operation span abs |
|---|---|---|---|
| Digit span bw max | $r$ = 0.46, $p$ = 0.000 | $r$ = 0.19, $p$ = 0.103 | $r$ = 0.17, $p$ = 0.138 |
| Digit span fw max | | $r$ = 0.32, $p$ = 0.005 | $r$ = 0.22, $p$ = 0.062 |
| Listening span | | | $r$ = -0.04, $p$ = 0.713 |

Note. Bw: backward maximum; fw: forward maximum; abs: absolute score.

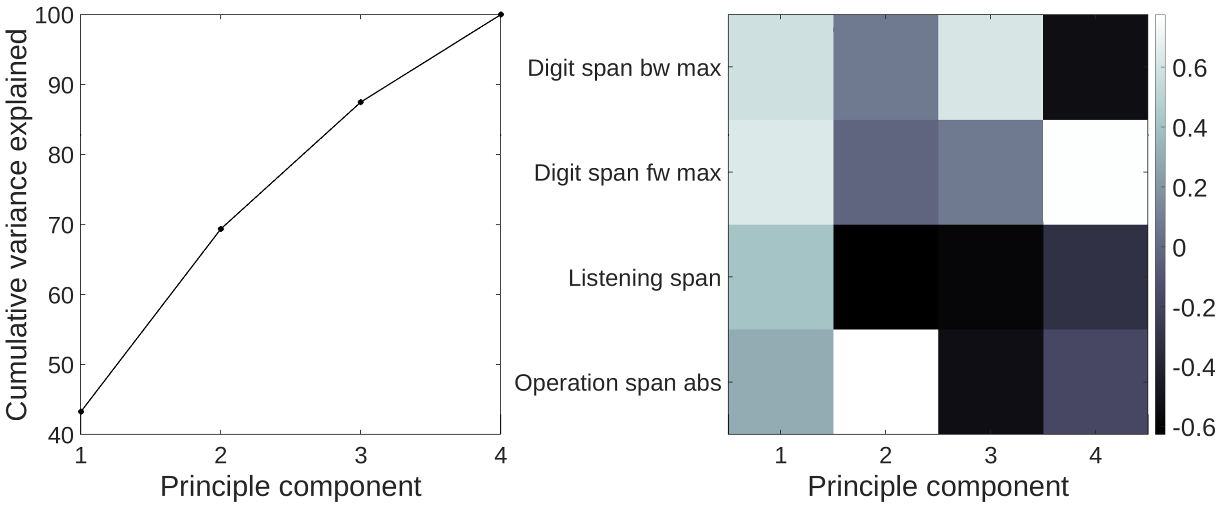


Figure S6a. Results for principal component analysis across three working memory tasks (digit span, listening span, and operation span). Cumulative variance explained (left panel) and factor loadings (right panel). Bw: backward maximum; fw: forward maximum; abs: absolute score.

### S6b Principal component analysis across all individual difference measures

The putative dopamine proxy variables spontaneous eye blink rate (sEBR), working memory capacity and impulsivity (BIS-15 score) were condensed to a single principal component using principal component analysis as implemented in MATLAB's (The MathWorks, Inc.) pca function. All proxy measures were *z*-scored before entering the analysis. The three variables were only weakly correlated with each other ($r_{sEBR – BIS-15}$ = 0.16, $p$ = 0.180; $r_{sEBR – WM}$ = -0.13, $p$ = 0.260; $r_{BIS-15 – WM}$ = -0.08, $p$ = 0.469). The cumulative variance explained and factor loadings are depicted in Figure S6b.

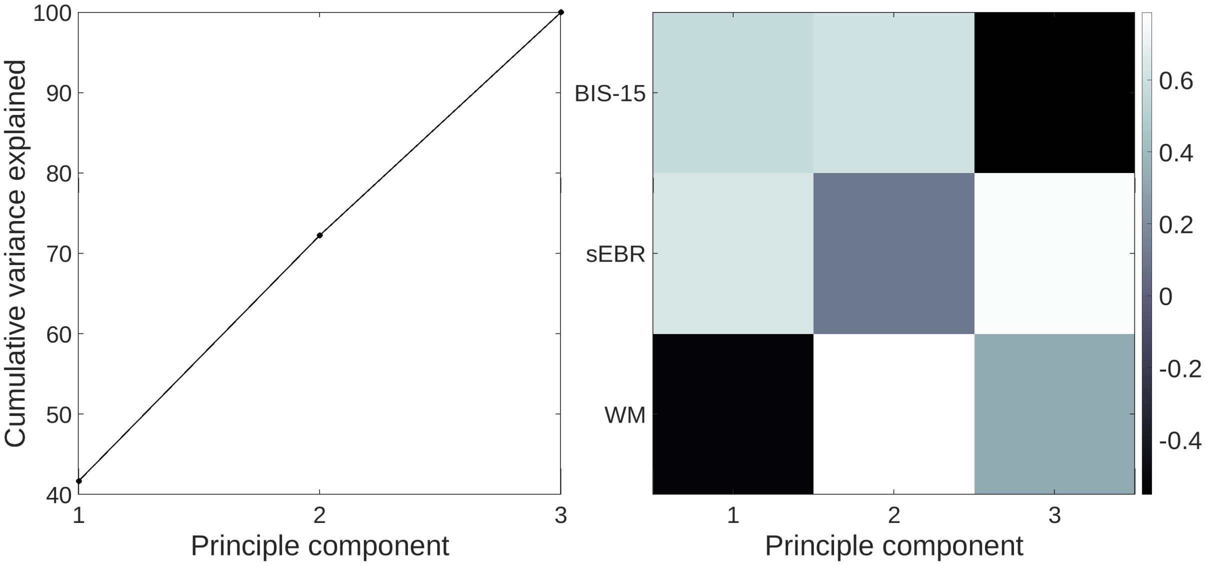


Figure S6b. Results for principal component analysis of spontaneous eye blink rate (sEBR), working memory capacity (WM) and impulsivity (BIS-15 score). Cumulative variance explained (left panel) and factor loadings (right panel).

### S6c Bayesian regression linear coefficients

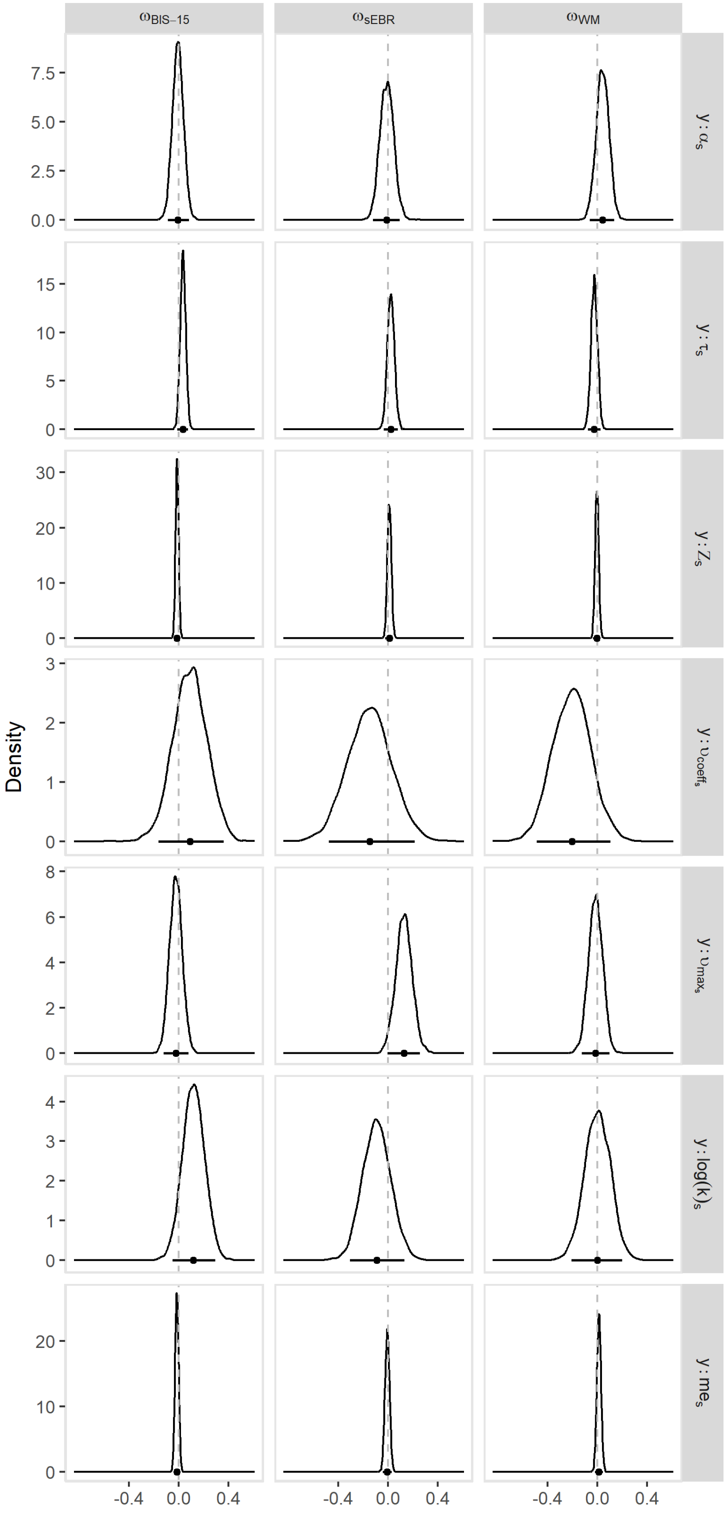


Figure S6c. Posterior distributions of the linear regression coefficients (ʊ) for the dopamine proxies impulsivity (BIS-15 scores), spontaneous eye blink rate (sEBR) and working memory capacity (WM), regressing the proxies onto the drug-shift (s) parameters of the drug-effect $DDM_{sigmoid}$. The posterior distribution of the intercept is not depicted. The horizontal solid line indicates the 95% highest posterior density interval, the vertical dashed line indicates x = 0. $\alpha$: boundary separation; $\tau$: non-decision time; $z$: starting-point bias; $\upsilon_{coeff}$: drift rate scaling factor; $\upsilon_{max}$: asymptote for $\upsilon_{coeff}$; $\log(k)_s$: drug effect for $\log(k)$; $me_s$: drug effect for $me$.

### S6d Bayesian regression quadratic coefficients

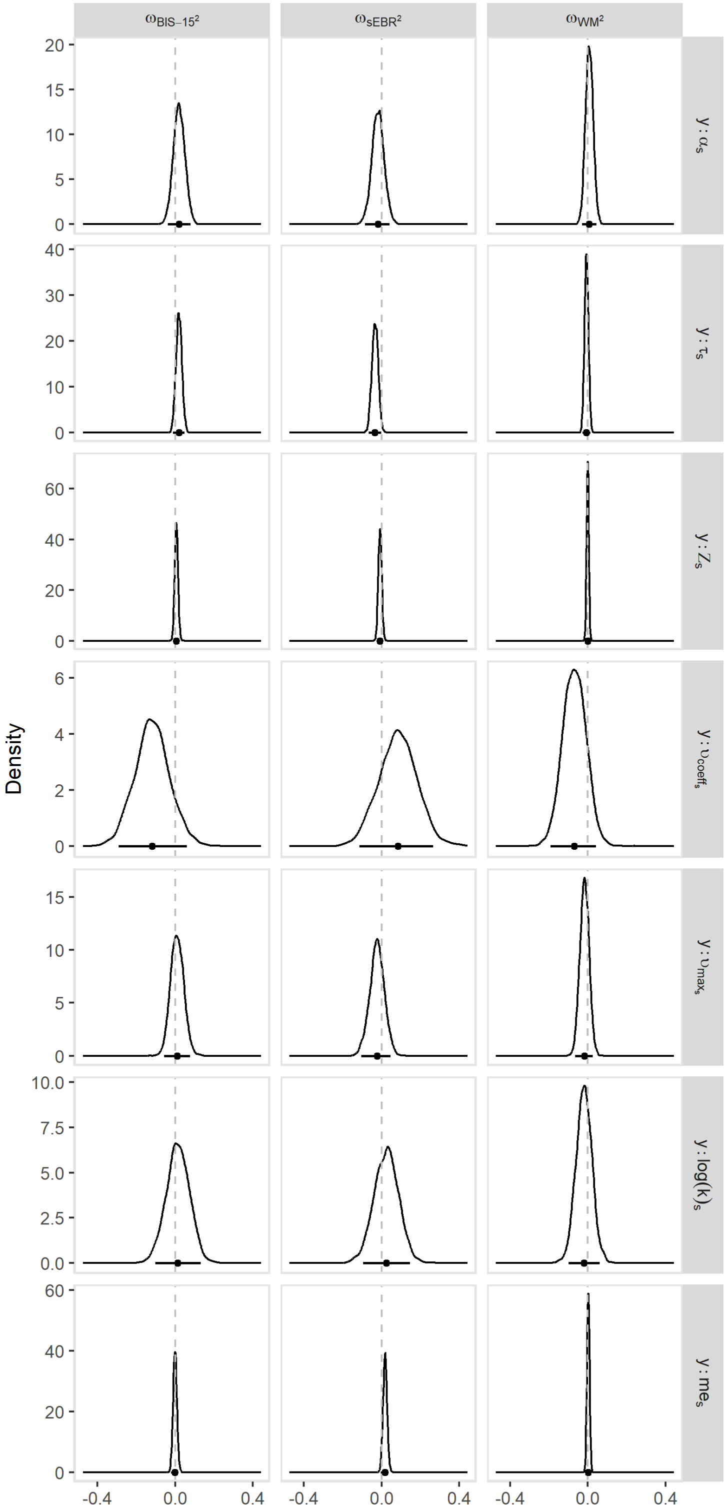


Figure S6d. Posterior distributions of the quadratic regression coefficients (ʊ) for the dopamine proxies impulsivity (BIS-15 scores), spontaneous eye blink rate (sEBR) and working memory capacity (WM), regressing the proxies onto the drug-shift (s) parameters of the drug-effect DDM-$_{sigmoid}$. The posterior distribution of the intercept is not depicted. The horizontal solid line indicates the 95% highest posterior density interval, the vertical dashed line indicates x = 0. $\alpha$: boundary separation; $\tau$: non-decision time; $z$: starting-point bias; $\upsilon_{coeff}$: drift rate scaling factor; $\upsilon_{max}$: asymptote for $\upsilon_{coeff}$; $\log(k)_s$: drug effect for $\log(k)$; $me_s$: drug effect for $me$.

Table S6e. Means and 95% HDIs for the posterior distributions of the linear (top) and quadratic (bottom) regression coefficients, regressing the putative dopamine proxies onto the drug-shift (s) parameters from the drug-effect $DDM_{sigmoid}$.

| | $\mho_{BIS-15}$ | | $\mho_{sEBR}$ | | $\mho_{WM}$ | |
|---|---|---|---|---|---|---|
| | *M* | *95% HDI* | *M* | *95% HDI* | *M* | *95% HDI* |
| $\alpha_s$ | -0.01 | [-0.09, 0.08] | -0.01 | [-0.12, 0.09] | 0.04 | [-0.06, 0.13] |
| $\tau_s$ | 0.03 | [-0.01, 0.08] | 0.02 | [-0.03, 0.08] | -0.03 | [-0.07, 0.03] |
| $z_s$ | -0.01 | [-0.04, 0.01] | 0.01 | [-0.02, 0.04] | 0.00 | [-0.03, 0.02] |
| $\upsilon_{coeff_s}$ | 0.09 | [-0.16, 0.36] | -0.15 | [-0.50, 0.19] | -0.21 | [-0.49, 0.10] |
| $\upsilon_{max_s}$ | -0.02 | [-0.12, 0.08] | 0.13 | [-0.01, 0.25] | -0.02 | [-0.12, 0.10] |
| $\log(k)_s$ | 0.12 | [-0.05, 0.29] | -0.09 | [-0.30, 0.14] | 0.00 | [-0.21, 0.20] |
| $me_s$ | -0.01 | [-0.04, 0.01] | -0.01 | [-0.04, 0.03] | 0.01 | [-0.02, 0.04] |
| | $\mho_{BIS-15}^2$ | | $\mho_{sEBR}^2$ | | $\mho_{WM}^2$ | |
| | *M* | *95% HDI* | *M* | *95% HDI* | *M* | *95% HDI* |
| $\alpha_s$ | 0.02 | [-0.04, 0.08] | -0.02 | [-0.08, 0.04] | 0.01 | [-0.03, 0.05] |
| $\tau_s$ | 0.02 | [-0.01, 0.05] | -0.03 | [-0.07, -0.00] | -0.01 | [-0.03, 0.01] |
| $z_s$ | 0.01 | [-0.01, 0.02] | -0.01 | [-0.03, 0.01] | 0.00 | [-0.01, 0.01] |
| $\upsilon_{coeff_s}$ | -0.12 | [-0.29, 0.06] | 0.08 | [-0.11, 0.27] | -0.07 | [-0.19, 0.04] |
| $\upsilon_{max_s}$ | 0.01 | [-0.06, 0.08] | -0.02 | [-0.10, 0.05] | -0.02 | [-0.07. 0.03] |
| $\log(k)_s$ | 0.01 | [-0.12, 0.13] | 0.02 | [-0.10, 0.14] | -0.02 | [-0.10, 0.06] |
| $me_s$ | 0.00 | [-0.02, 0.02] | 0.02 | [-0.00, 0.04] | 0.00 | [-0.01, 0.01] |

Note. Values for the posterior distribution of the intercept are not listed. DDM: drift diffusion model; BIS-15: Baratt Impulsiveness Scale; sEBR: spontaneous eye blink rate; WM: working memory capacity; HDI: highest posterior density interval; $\alpha$: boundary separation; $\tau$: non-decision time; $z$: starting-point bias; $\upsilon_{coeff}$: drift rate scaling factor; $\upsilon_{max}$: asymptote for $\upsilon_{coeff}$; $\log(k)_s$: drug effect for $\log(k)$; $me_s$: drug effect for $me$.

### S6e Relationship between drug effects and DA proxy measures

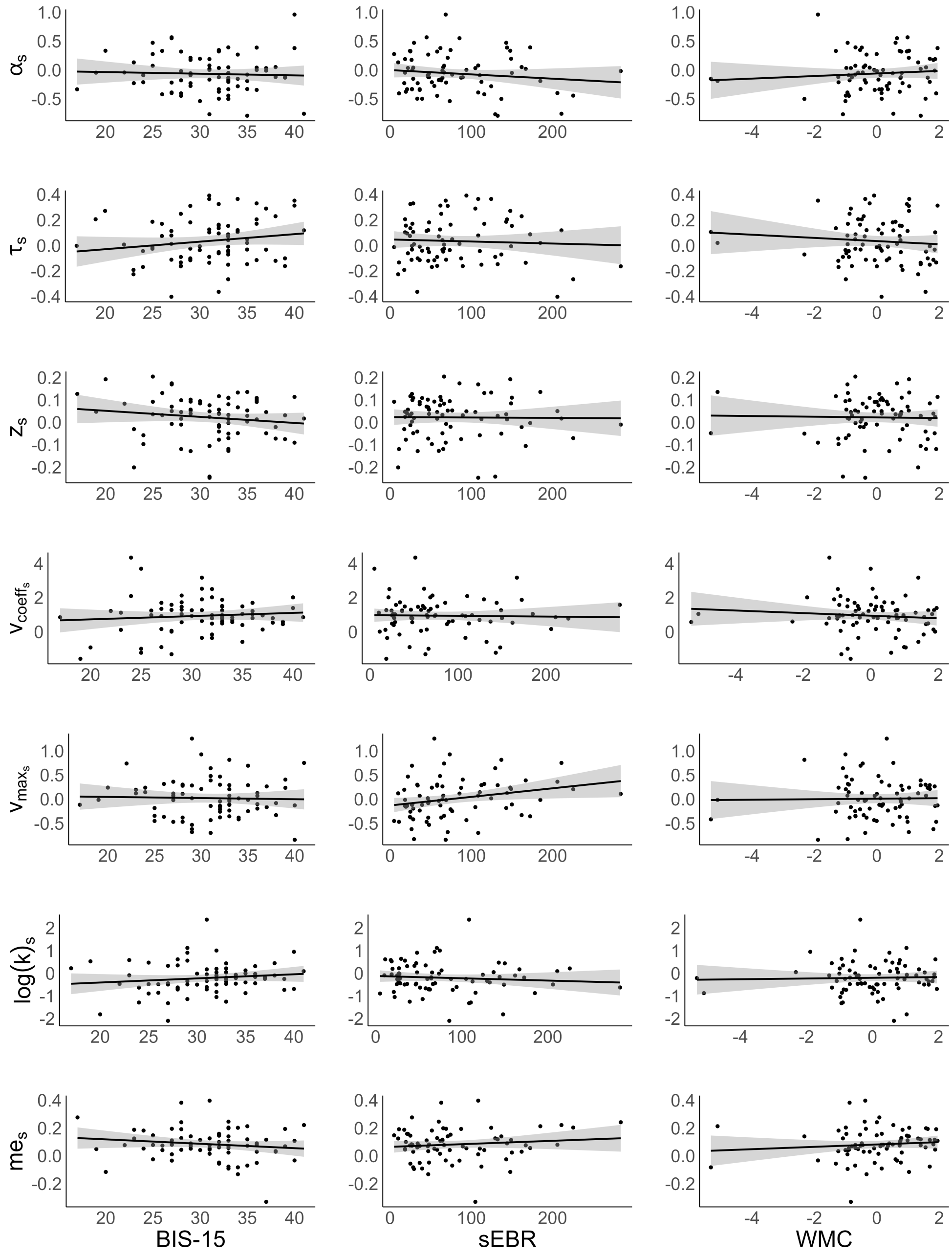


Figure S6e. Relationship between the putative dopamine proxy measures impulsivity (BIS-15 scores), spontaneous eye blink rate (sEBR) and working memory capacity (WMC) and the drug effects (means of single-subject posterior distributions for the drug effect parameters of the drug-effect $DDM_{sigmoid}$. Points: single-subject values; solid black line: regression line; grey ribbon: standard error. $s$: drug effect (additive "shift") parameter; α: boundary separation; τ: non-decision time; $z$: starting-point bias; $v_{coeff}$: drift rate scaling factor; $v_{max}$: asymptote for $v_{coeff}$; log(k): discount rate; me: magnitude effect.
